# Myocarditis in naturally infected pets with the British variant of COVID-19

**DOI:** 10.1101/2021.03.18.435945

**Authors:** Luca Ferasin, Matthieu Fritz, Heidi Ferasin, Pierre Becquart, Vincent Legros, Eric M. Leroy

## Abstract

Domestic pets can contract SARS-CoV-2 infection but, based on the limited information available to date, it is unknown whether the new British B.1.1.7 variant can more easily infect certain animal species or increase the possibility of human-to-animal transmission. In this study, we report the first cases of infection of domestic cats and dogs by the British B.1.1.7 variant of SARS-CoV-2 diagnosed at a specialist veterinary hospital in the South-East of England. Furthermore, we discovered that many owners and handlers of these pets had developed Covid-19 respiratory symptoms 3-6 weeks before their pets became ill and had also tested PCR positive for Covid-19. Interestingly, all these B.1.1.7 infected pets developed atypical clinical manifestations, including severe cardiac abnormalities secondary to myocarditis and a profound impairment of the general health status of the patient but without any primary respiratory signs. Together, our findings demonstrate for the first time the ability for companion animals to be infected by the B.1.1.7 variant of SARS-CoV-2 and raise questions regarding its pathogenicity in these animals. Moreover, given the enhanced infectivity and transmissibility of B.1.1.7 variant for humans, these findings also highlights more than ever the risk that companion animals may potentially play a significant role in SARS-CoV-2 outbreak dynamics than previously appreciated.

## Introduction

The COVID-19 pandemic secondary to severe acute respiratory syndrome coronavirus 2 (SARS-CoV-2) variant, carrying the Spike (S) protein amino acid change D614G (referred to as B. 1 variant), has encompassed several millions of cases around the world. This global situation has favored the appearance of numerous genomic mutations, some of which have generated variants having selective advantages^1^. Three notable variants have emerged in late fall 2020 in several countries, which then spread rapidly across the world, including B.1.1.7 (also referred to as 20I/N501Y.V1) first detected in England ^2^, B.1.351 (20J/N501Y.V2) first detected in South Africa and the recently identified “Brazil” variant P.1 (20I/N501Y.V3). These three variants carry a constellation of genetic mutations, including those at the level of the S protein receptor-binding domain (RDB), which is essential for binding to the cell host ACE-2 receptor to facilitate virus entry.

The B.1.1.7 variant, also referred to as variant of concern (VOC) 202012/01 or 20I/501Y.V1, is estimated to have emerged in September 2020 in Kent, a county in the South-East of England, and has rapidly outcompeted pre-existing variants in England as the consequence of an increased transmissibility and infectivity ^2^. Multiple lines of evidence indicate that its enhanced transmissibility is driven by the N501Y mutation and the amino acid Δ69/70 deletion in RDB ^3^. Consequently, the incidence of B.1.1.7 increased rapidly during a national lockdown implemented by the Government of the United Kingdom from 5 November to 2 December 2020, despite rigorous restrictions, causing an extraordinary surge of COVID-19 cases particularly affecting the Greater London area. As of 7 February 2021, VOC 202012/01 comprised roughly 95% of new SARS-CoV-2 infections in the United Kingdom and has now been identified in at least 86 countries.

Several cases of SARS-CoV-2 infection have also been reported worldwide in domestic pets (especially cats and dogs) and it has been suggested that these animals became infected by their owners or handlers. Infections of domestic pets mostly resulted in unapparent to mild digestive and respiratory symptoms such as cough, runny nose, sneezing and conjunctivitis ^4–6^. Intriguingly, despite the uncontrolled surge of COVID-19 cases occurring in the UK since November 2020, no infections by SARS-CoV-2 have been reported in companion animals so far. More surprisingly, to the best of the authors’ knowledge, natural infection of any animal by the B.1.1.7 variant has never been documented, neither in England nor anywhere else.

## Results

We report a sudden increased number of domestic dogs and cats presented with myocarditis at the Cardiology Department of The Ralph Veterinary Referral Centre (RVRC), based on the outskirts of London (UK), between December 2020 and February 2021, with an unexpected rise in incidence from 1.4% to 12.8% (8.5% in cats and 4.3% in dogs). This sudden surge of cases appeared to mimic the curve and timeline of the COVID-19 human pandemic in the UK due to the B.1.1.7 variant, starting in mid-December 2020, peaking at the end of January 2021, before returning to the historical rate by mid-February 2021.

None of these patients with myocarditis had a previous history of heart disease and their clinical presentation was similar and characterized by acute onset of lethargy, inappetence, tachypnea/dyspnea (secondary to the presence of congestive heart failure), and, in some cases, syncopal events. Diagnostic investigations revealed the presence of elevated cardiac troponin-I (median 6.8; range 0.68 to 61.1 ng/mL [normal reference range 0.0-0.2 ng/mL]) accompanied by echocardiographic evidence of myocardial remodeling and/or signs of pleural effusion and/or pulmonary edema, often confirmed on thoracic radiographs and/or severe ventricular arrhythmias on electrocardiography (see Supplementary Figure S1). All affected animals made a remarkable improvement with cage rest, oxygen therapy, acute diuresis and, in some cases, anti-arrhythmic therapy with sotalol and fish oil supplementation before being discharged on oral medications after a few days of intensive care. Notably, most owners and handlers of these pets with myocarditis had developed Covid-19 respiratory symptoms within 3-6 weeks before their pets became ill and many of these owners had tested PCR positive for Covid-19. Given this coincidence and the intriguing simultaneous evolution of myocarditis in these pets and the B1.1.7 COVID-19 outbreak in UK, we decided to investigate SARS-CoV-2 infection in these animals. For this purpose, serum samples as well as oro/nasopharyngeal and rectal swabs were collected from seven animals (six cats and one dog) at initial presentation at the RVRC between January 22 and February 10, 2021 (Table 1 and Supplementary Table S1). During the same period, we collected blood samples from four other pets (two cats and two dogs) during their recovery, 2-6 weeks after they developed signs of myocarditis. None of the 11 animals with myocarditis developed any influenza-like symptoms and they all clinically improved within a few days of intensive care, although one cat (LL) represented one week after discharge with a relapse of her clinical signs, characterized by profound lethargy and uncontrolled ventricular tachycardia, prompting her owners to elect for euthanasia. All cats and dogs were neutered and aged between one to 12 years. Following collection, all samples were stored at −20°C until transportation in ice to MIVEGEC laboratory at Montpellier, France, for serological and virological investigations.

**Table 1.** Characteristics of dogs and cats diagnosed with myocarditis at The Ralph Veterinary Referral Centre between January 22 and February 10, 2021

|  | Species | Breed | Age | Sex | Days | General symptoms | Cardiac abnormalities | Troponin ng/mL | Covid-19+ contact | SARS-CoV-2 ddPCR | Serology |
| --- | --- | --- | --- | --- | --- | --- | --- | --- | --- | --- | --- |
| **BBK** | cat | DSH | 9 | M | 2 | Lethargy<br>Inappetance | CHF | 7.9 | Yes | - | - |
| **HY** | cat | DSH | 9 | M | 2 | Lethargy<br>Inappetance | CHF, VA | 0.68 | Yes | 33 copies RNA/μL | - |
| **CH** | cat | Manx | 12 | F | 2 | Lethargy | CHF, VA | 6.8 | Unknown | - | - |
| **LL** | cat | Sphynx | 10 | F | 3 | Syncope | CHF, VA | 45.6 | Unknown | 12 copies RNA/μL | - |
| **MR** | dog | Labrador | 9 | F | 4 | Lethargy, Inappetance<br>Hemorrhagic diarrhea | VA | 43.5 | Yes | 13 copies RNA/μL | - |
| **DB** | cat | DSH | 9 | M | 8 | Lethargy | CHF | 1.31 | Unknown | - | + |
| **FB** | cat | Scottish Fold | 1 | M | 10 | Lethargy<br>Inappetance | CHF | 12.1 | Unknown | - | - |
| **DP** | dog | Mastiff | 8 | F | 14 | Syncope | CHF, VA | 2.5 | Unknown | NA | - |
| **SC** | cat | Siberian | 1 | F | 28 | Lethargy | CHF | 4.92 | Yes | NA | + |
| **KEO** | dog | Dalmatian | 8 | M | 37 | Syncope | VA | 61.1 | Yes | NA | + |
| **OR** | cat | Persian | 1 | M | 64 | Lethargy | CHF, VA | 0.83 | Unknown | NA | - |

Days: days between disease onset and the medical consultation at the clinics, including sampling.
DSH: Domestic shorthair cat. CHF: Congestive heart failure. VA: Ventricular arrhythmia. NA: Not available.

Oro/nasopharyngeal and rectal swabs were tested using the droplet digital RT-PCR (ddPCR) targeting one region specific to the SARS-CoV-2 N gene and two regions of the Spike protein gene specific to the three current predominant SARS-CoV-2 variants, namely 20I/N501Y.V1, 20J/N501Y.V2 and 20I/N501Y.V3. One target region, containing the N501Y mutation, is common to the three variants and the other target region, containing the Δ69-70 deletion, is specific to the B.1.1.7 variant. Sera were tested for SARS-CoV-2-specific IgG using three microsphere immunoassays (MIA) detecting IgG binding to the N protein, the S1-RBD-protein or the S trimeric protein, as well as a retrovirus-based pseudoparticle assay detecting SARS-CoV-2 neutralizing antibodies (see Materials and methods).

All oro/nasopharyngeal swabs were found SARS-CoV-2 ddPCR negative. However, ddPCR positive signals were obtained for the three regions from the rectal swabs from three of seven animals (two cats and one dog), indicating infection with the British B.1.1.7 variant. The RNA concentration ranged from 12 to 33 copies/μL of specimen, indicating low viral load (Table 1). In addition, one animal sampled during the acute phase of the disease which tested ddPCR negative, as well as two of four animals sampled during the recovery period, were found to have SARS-CoV-2 antibodies. Therefore, in total, six of our 11 investigated animals were shown SARS-CoV-2 positive, either by ddPCR or by serology. More interestingly, considering only the five animals from which owners or handlers were laboratory confirmed Covid-19 positive, four were shown SARS-CoV-2 positive (Table 1).

## Discussion

To our knowledge, this is the first report of infection of both cats and dogs by the British B.1.1.7 variant of SARS-CoV-2. Given the enhanced infectivity and transmissibility of B.1.1.7 variant for humans, the discovery of B.1.1.7 infected cats and dogs highlights more than ever the risk that companion animals may potentially play a significant role in SARS-CoV-2 outbreak dynamics than previously appreciated. Further studies are therefore urgently required to investigate the likelihood of pet-to-pet transmission, as well as pet-to-human transmission of the B.1.1.7 variant and to show whether the N501Y mutation and the Δ69-70 deletion render the virus more infectious for these animals.

The other remarkable and unexpected finding of our study is the development of unusual clinical manifestations in B.1.1.7 infected cats and dogs, including severe cardiac abnormalities secondary to myocarditis and a profound impairment of the general health status of the patient but without any primary respiratory signs. With the exception of only one cat from Spain, which developed cardiorespiratory failure resulting in severe respiratory distress ^7^, both natural and experimental SARS-CoV-2 infections of cats and dogs have so far been reported to be either asymptomatic or display mild upper respiratory disease. Although that the B.1.1.7 infection in humans seems to be associated with higher COVID-19 mortality or clinical severity, the association between myocarditis and B.1.1.7 infection in domestic pets has to be acknowledged and addressed ^8^. In this context, it is important to highlight the fact that myocarditis associated with multisystem inflammatory syndrome is also a well-recognized complication of COVID-19 in people (both adults and children) probably from exaggerated immune response of the host ^9,10.^

Together, our findings demonstrate the ability for companion animals to be infected by the B.1.1.7 variant of SARS-CoV-2 and raise questions regarding its pathogenicity in these animals. Therefore, there is an urgent need to greatly accelerate and strengthen the investigations and surveillance of animal infections by highly-transmissible variants such as British B.1.1.7, South-African B1.351 and Brazilian P.1 variants as part of the global response to the ongoing multi-species COVID-19 pandemic.

## Materials and methods

### RNA extraction

Rectal and oro/naso-pharyngeal swabs were resuspended by vortexing in 300 μL of PBS. Total RNA was extracted from 200 μL of supernatant of rectal swab and from 200 μL of viral transport medium of nasopharyngeal swabs. Extraction was performed on the extraction system IndiMag 48 (Indical Bioscience), using magnetic bead technology, with the IndiMag Pathogen kit according to the manufacturer’s instructions. The elution volume was 100 μL.

### One step dRT-PCR

The RT-dPCR procedure was performed following the manufacturer’s instructions using the QIAcuity 8, 5-plex (Qiagen, Germany), the QIAcuity One-Step Viral RT-PCR Kit (Cat No. 1123145, Qiagen, Germany) and the 24-well 26K Nanoplates (Cat No. 250001, Qiagen, Germany). The ddRT-PCR technique showed higher sensitivity and specificity compared to RT-qPCR for diagnosis of COVID-1911.

Briefly, the RT-dPCR reaction mixture was assembled as follows: 4x One-Step Viral RT-PCR Master Mix 10μl, 100x Multiplex Reverse Transcription Mix 0.4μl, 20x of set of primers and probes 0149, 0130, 0150 (ref IAGE) 2μl x3 (6μL), RNase free water 22.6μl, and RNA template 1 μl, in a final volume of 40 μl. 0130 target 2019-nCoV_N2 region NC_045512v2 fluorophore HEX, amplicon length 67bp. 0149 target S region: mutation deletion 69-70, lineage B1.1 & B1.258, fluorophore HEX, amplicon length 100bp. 0150 target S region: mutation N501Y, lineage B1.1.7, fluorophore Cy5, amplicon length 133bp.The sequences are confidential and are filed under the number EP20306715.2 The mixture was prepared in a pre-plate and then transferred into the 24-well 26K Nanoplate. The later was then loaded to the QIAcuity 8 instrument, which is a fully automated system. The workflow included i) priming and rolling step in order to generate and isolate the chamber partitions, ii) the amplification step under the following cycling protocol: 50 °C for 40 min for reverse transcription, 95 °C for 2 min for enzyme activation, 95 °C for 5 s for denaturation and 60 °C for 30 s for annealing/extension for 40 cycles, and iii) the imaging step was done by reading in the following channels FAM, HEX, and CY5. The full workflow time was around 2 hours for the three steps. The experiments were performed using a negative control (no template control, NTC) and a positive control (a patient’s sample confirmed positive by RT-PCR with our routine diagnostic testing). All reactions had at least 25,400 partitions. Data were analysed using the QIAcuity Suite Software V1.1.3 (Qiagen, Germany) and expressed as copies/μl.

### Microsphere immunoassay

Cat and dog serum samples were tested using a multiplex Microsphere immunoassay (MIA). 10μg of three recombinant SARS-CoV-2 antigens: nucleoprotein (N), receptor binding domain (RBD) and trimeric spike (tri-S) were used to capture specific serum antibodies. Distinct MagPlex microsphere sets (Luminex Corp) were respectively coupled to viral antigens using the amine coupling kit (Bio-Rad Laboratories) according to manufacturers’ instructions, whereas a microsphere set coupled with recombinant human protein (O6-methylguanine DNA methyltransferase) was used as control in the assay. The MIA procedure was performed as described previously ^12^. Briefly, microsphere mixtures were successively incubated protected from the light on an orbital shaker with serum samples (1:400), biotinylated protein A and biotinylated protein G (4 μg/ml each) (Thermo Fisher Scientific) and Streptavidin-R-Phycoerythrin (4 μg/ml) (Life technologies). Measurements were performed using a Magpix instrument (Luminex). Relative Fluorescence Intensities (RFI) were calculated for each sample by dividing the MFI signal measured for the antigen-coated microsphere sets by the MFI signal obtained for the control microsphere set, to account for nonspecific binding of antibodies to beads. Specific seropositivity cut-off values for each antigen were set at three standard deviations above the mean RFI of the 29 dogs and 30 cats serum samples sampled before 2019. Based on the pre-pandemic population, MIA specificity was set at 96.6% for N protein for dogs and cats, at 96.6% for RBD for cats and 100% for dogs and 100% for tri-S for cats and 96.6% for dogs.

### Neutralization activity measurement

To measure the neutralizing activity in cat and dog sera, a MLV-based pseudoparticle carrying a GFP reporter pseudotyped with SARS-CoV2 spike (SARS-CoV-2pp) was used. Each SARS-CoV2 positive sample detected by MIA was processed according to neutralization procedure as previously described ^13^. The level of infectivity was expressed as % of GFP positive cells and compared to cells infected with SARS-CoV-2pp incubated without serum. Pre-pandemic sera from France was used as negative controls, and anti-SARS-CoV-2 RBD antibody was used as positive control.

## Acknowledgements

We are grateful to the pet owners for giving us their permission to sample their pets. We also thank Dr Laurent Locquet and Dr Altin Cala for contributing to the clinical management of these patients. Thanks also to Franz Durandet (IAGE – Ingénierie et Analyse en Génétique Environnementale, Grabels, France), Alix de Mont-Marin (Innovative Diagnostics, France) and Afif M. Abdel Nour (Qiagen, France) for their help in molecular biology. We also thank François Loïc Cosset, Solène Denolly, Bertrand Boson from CIRI – Centre International de Recherche en Infectiologie, Team EVIR, Univ Lyon, Université Claude Bernard Lyon 1, Inserm, U111, UMR5308, ENS Lyon, for the development of the seroneutralization technique. Finally, we thank Dr Thierry Buronfosse for the kind gift of pre-pandemic sera.

