## Supplementary Figure S1 for "Myocarditis in naturally infected pets with the British variant of COVID-19"

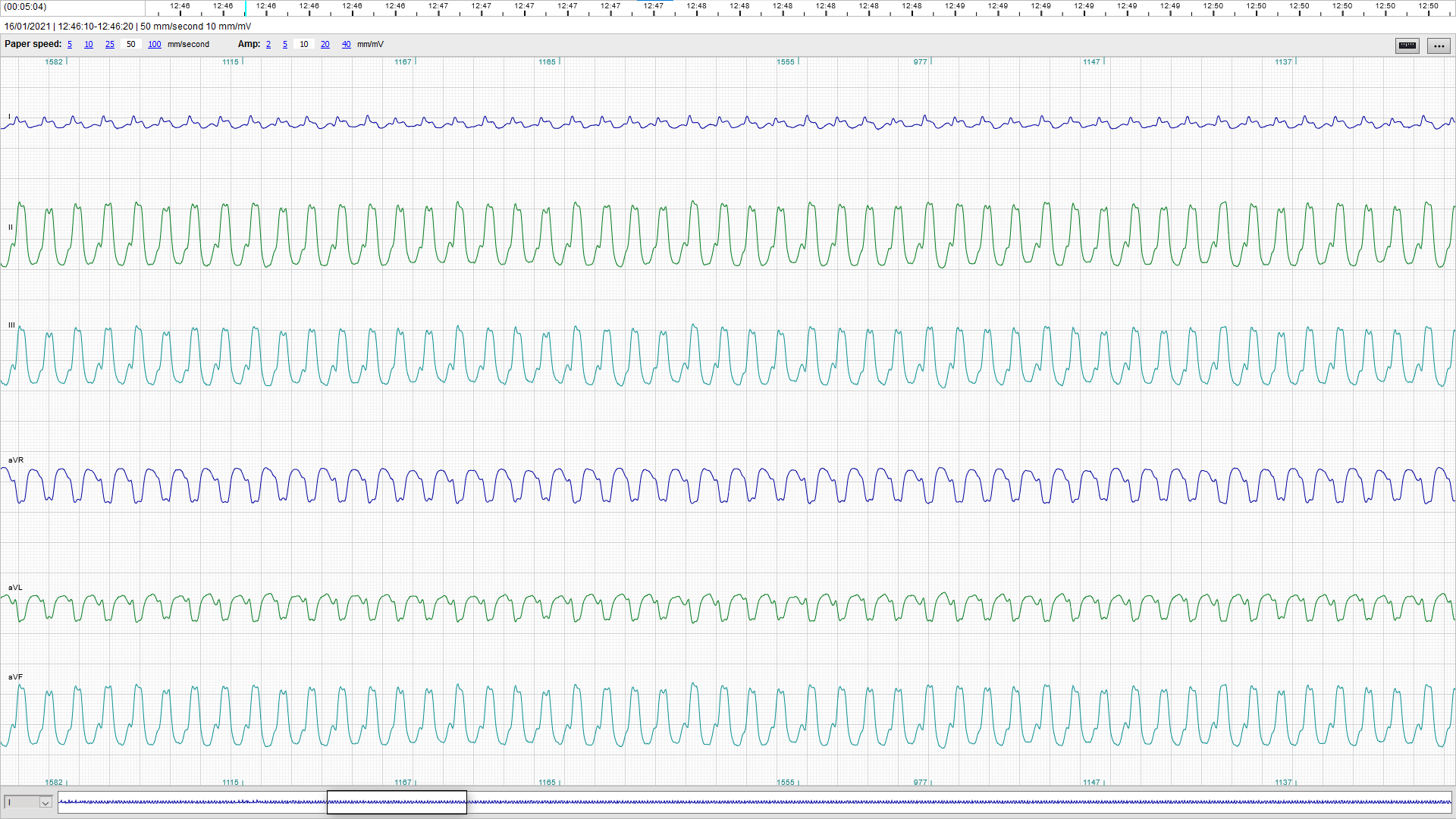
**Supplementary Figure 1**: ECG trace recorded from one of the dogs presented with signs of acute myocarditis showing sustained monomorphic ventricular tachycardia at 320 bpm (sample obtained from a 5-minute continuous 6-lead ECG recording, 50 mm/s, 10 mm/mV).
