## Supplementary Table S1 for "Myocarditis in naturally infected pets with the British variant of COVID-19"

**Supplementary Table 1. Characteristics of dogs and cats diagnosed with myocarditis at The Ralph Veterinary Referral Centre between January 22 and February 10, 2021**

|  | Species | Breed | Age | Sex | Days | General symptoms | T, °C | Cardiac  abnormalities | Troponin  ng/mL | Covid-19+ contact | Serology  N protein | Serology  RBD | Serology  S trimeric | SN | SARS-CoV-2  PCR (rectal) | SARS-CoV-2  PCR (nasoph) |
| --- | --- | --- | --- | --- | --- | --- | --- | --- | --- | --- | --- | --- | --- | --- | --- | --- |
| BBK | cat | DSH | 9 | M | 2 | Lethargy  Inappetance | 37.1 | CHF | 7.9 | Yes | - | - | - | - | - | - |
| HY | cat | DSH | 9 | M | 2 | Lethargy  Inappetance | 37.0 | CHF, VA | 0.68 | Yes | - | - | - | - | 33 copies RNA/µL | - |
| CH | cat | Manx | 12 | F | 2 | Lethargy | 35.2 | CHF, VA | 6.8 | Unknown | - | - | - | - | - | - |
| LL | cat | Sphynx | 10 | F | 3 | Syncope | 37.9 | CHF, VA | 45.6 | Unknown | - | - | - | - | 12 copies RNA/µL | - |
| MR | dog | Labrador | 9 | F | 4 | Lethargy, Inappetance  Hemorrhagic diarrhea | 37.6 | VA | 43.5 | Yes | - | - | - | - | 13 copies RNA/µL | - |
| DB | cat | DSH | 9 | M | 8 | Lethargy | 37.8 | CHF | 1.31 | Unknown | + | - | - | - | - | - |
| FB | cat | Scottish Fold | 1 | M | 10 | Lethargy  Inappetance | 37.3 | CHF | 12.1 | Unknown | - | - | - | - | - | - |
| DP | dog | Mastiff | 8 | F | 14 | Syncope | 38.0 | CHF, VA | 2.5 | Unknown | - | - | - | - | NA | NA |
| SC | cat | Siberian | 1 | F | 28 | Lethargy | 38.5 | CHF | 4.92 | Yes | + | + | + | - | NA | NA |
| KEO | dog | Dalmatian | 8 | M | 37 | Syncope | 38.2 | VA | 61.1 | Yes |  | + | + | + | NA | NA |
| OR | cat | Persian | 1 | M | 64 | Lethargy | 37.5 | CHF, VA | 0.83 | Unknown | - | - | - | - | NA | NA |
